# Serum proteomics in newborns reveals a multi-pathway molecular panel for neonatal sepsis, encompassing pathogen sensing, leukocyte recruitment, and endothelial injury

**DOI:** 10.64898/2026.05.27.728097

**Authors:** Kevin Roger, Najah Fatou Coly, Ines Metatla, Fatou Diallo Agne, Idrissa Basse, Papa Madiéye Gueye, Cerina Chhuon, Ida Chiara Guerrera

## Abstract

**Background:** Neonatal sepsis is a major cause of morbidity and mortality, particularly in low- and middle-income countries such as Senegal, where incidence is 78–104 per 1,000 live births and mortality exceeds 20 per 1,000, with case fatality rates around 36%. Diagnosis is difficult due to non-specific clinical signs and lack of molecular biomarkers, highlighting the need for improved early diagnostic molecular panel that could be applied even outside hospital settings.

**Objectives:** Compare neonatal and adult serum proteomes to establish a reference and identify serum protein biomarkers of neonatal sepsis.

**Methods:** Serum samples from Senegalese neonates and adults were analyzed using data-independent acquisition (DIA) proteomics on neat serum (Evosep-timsTOF HT platform). The cohort comprised 22 neonates with confirmed sepsis (CS), 6 neonates with non-confirmed sepsis (NCS), 17 healthy newborn controls (HC), 6 unclassified, 9 outliers and 20 healthy adults. Downstream analyses included differential protein abundance testing, unsupervised clustering, weighted gene co-expression network analysis (WGCNA), and correlation analyses with clinical parameters.

**Results:** We identified 979±20 proteins in newborns versus 718±40 in adults. Newborns showed reduced immune-response related proteins, a narrower dynamic range, and increased structural proteins such as collagens, consistent with immune immaturity and tissue development. Unsupervised WGCNA analysis led to a 53-protein cluster discriminated CS from NCS/HC. Some of these dysregulated proteins identified have already been reported in independent studies using different approaches in neonatal and/or adult sepsis. Our larger panel however of identified markers maps to three major biological processes involved in sepsis: (i) pathogen sensing (LBP, CD14), and acute-phase inflammation (e.g. CRP, SAA1/2, ORM1/2); (ii) innate immune activation and leukocyte recruitment (e.g., FCGR3A, CSF1R, CD163, CD206) and final platelet exhaustion, (e.g., PF4, PPBP, THBS1, GP5); (iii) endothelial injury and microvascular dysfunction with tissue remodeling (e.g., ICAM1, VCAM1, VWF, SPARC) with simultaneous loss of protective lipoproteins and serpins (e.g., APOA1, APOA2, APOM, SERPINA4, SERPINA5)

**Conclusion:** This study provides a very comprehensive neonatal serum proteome characterization and identifies a protein panel of proteins mapped to three major processes in sepsis.

## Introduction

Alongside prematurity, sepsis remains one of the main challenges in neonatal medicine, and they are responsible together for nearly 50% of deaths among children under the age of 5 worldwide^1–3^, reaching approximately 21–23% of neonatal deaths in some developing countries^4^. Indeed, the global burden is particularly severe in low- and middle-income countries such as Senegal, where community-based studies have reported incidences of severe neonatal bacterial infections ranging from 78 to 104 cases per 1,000 live births^4^, approaching case-fatality rates of 36%.

This is due to fragmented healthcare system, inequities in care, but mainly to the significant limitations of current diagnostic approaches. The first challenge is linked to the non-specific nature of neonatal sepsis symptoms—temperature instability, feeding difficulties, lethargy, and respiratory distress—all of which can easily be confused with other neonatal conditions. Secondly, blood culture, the gold standard for confirming bacteremia, requires 24–72 hours for results and has low sensitivity in neonates due to small sample volumes and prior maternal antibiotic exposure. Consequently, given the long delay and challenges to obtain reliable results from blood culture, clinicians rely on inflammatory biomarkers such as C-reactive protein (CRP), procalcitonin, and white blood cell counts to guide empirical antibiotic treatment decisions. However, classical markers such as CRP fail to reliably distinguish bacterial sepsis from sterile systemic inflammatory response syndrome (SIRS)5. Finally, inflammatory markers are not only non-specific to bacterial infection but also demonstrate considerable variability during the early neonatal period, when physiological developmental processes can mimic pathological inflammation^6,7^.These perinatal physiological changes could substantially confound the interpretation of results if inflammatory markers of adult sepsis are used for newborns cases. Global proteomics studies have been conducted to explore age-related remodeling of the plasma proteome in the first days of life. Early studies, as well as more extensive later proteomics analysis, have demonstrated that proteins such as fibrinogen chains, haptoglobin, clusterin, kininogen, and hemopexin are higher in adults, while α2-macroglobulin, complement C3, complement factor B, vitamin D–binding protein, heparin cofactor II, and bikunin are more abundant in neonates and children^8,9^. Moreover, longitudinal analysis during the first week of life shows dynamic changes in interferon signaling, Toll-like receptor pathways, complement activation, and neutrophil-associated signatures between day 0 and day 7 of life^6,7^. These studies have also documented a transient acute-phase response occurring at day 1 of life, characterized by sharp increases in acute-phase proteins such as haptoglobin and serum amyloid A, alongside concurrent decreases in negative acute-phase proteins like α2-macroglobulin and clusterin^6^.

Finally, the lack of consensus in the definition of neonatal sepsis makes it difficult to classify them even with a positive blood culture and hampers large scale association studies^10–12^. This diagnostic ambiguity, combined with the accelerated disease progression characteristic of neonatal sepsis, creates an urgent need for early and specific molecular markers that can distinguish true bacterial infection from other illness states in the critical first days of life.

MS-based serum/plasma proteomics, despite all its challenges, offers a compelling approach to address these gaps by providing an unbiased view of the neonatal landscape during infection and inflammation, as today can allow the quantification of over 1000 proteins^13^. Targeted immunoassay-based studies have been informative to quantify specific cytokine signatures associated with neonatal sepsis—such as elevations in IL-6, IL-10, TNF-α, and alterations in IFN-γ and their ratios^14^. However, these approaches are inherently limited to pre-selected panels and may miss broader protein signatures with diagnostic potential. Indeed, given that sepsis involves multiple biological processes, from both pathogen recognition, inflammatory response, immunoregulatory transition and vascular injury ^15,16^ a broader view of the proteome is essential to capture proteins involved in these processes and that could distinguish ongoing bacterial infection from transient tissue injury or sterile inflammation. In adult sepsis cohorts, some markers (CD64, procalcitonin, presepsin, sTREM-1, occasionally in multi-marker panels) show better diagnostic performance than CRP for sepsis vs non-infectious SIRS while LBP alone shows equal or worse, accuracy compared with CRP^17^. Beyond these proteins, numerous markers, including complement/coagulation factors, acute-phase reactants, adhesion molecules, and endothelial markers have also shown association with sepsis, death or severity^18,19^. However, many of these proteins also vary in newborns compared to adults’ serum. This motivated our choice to conduct an in-depth, age-specific proteomic investigation.

One recent publication reports the MS –based proteomics analysis of a large cohort of adults’ patients with confirmed sepsis and non-infectious systemic inflammatory. Despite the very limited coverage of 110 proteins quantified, they could point out to PPBP, VWF, both involved in endothelial/prothrombotic activation, as the most discriminating proteins between the two groups^19^. For newborn, a recent proteomic study on neonatal sepsis reports quantification of 255 proteins that were differentially abundant in cord blood of 14 infants with early onset sepsis (EOS) compared with 150 infants without EOS. Five proteins were significantly enriched in the plasma of infants with culture-confirmed EOS: CRP, lipopolysaccharide-binding protein (LBP), SAA1, leucine-rich α-2-glycoprotein 1 (LRG1), and serine proteinase inhibitor A3 (SERPINA3). These are all acute-phase reactant proteins that are upregulated in the plasma in response to inflammation^20^.

The present study aims to thoroughly characterize with unprecedented depth the neonatal serum proteome in the first days of life and to identify protein markers specific to the sepsis processes that can distinguish non-infectious systemic inflammation and confirmed bacterial sepsis in a Senegalese neonatal cohort.

## Methods

### Patients inclusion

The study was approved by the Research Ethics Committee of Cheikh Anta Diop University in Dakar. All mothers of recruited patients gave written informed consent. A total of 20 healthy adults, 20 healthy newborn and 40 newborns suspected of infection based on suggestive clinical symptomatology and anamnestic criteria were included in the study (Table 1, Table 2). Regarding the inclusion criteria, newborns suspected of having sepsis were included based on a combination of nonspecific clinical signs, maternal risk factors, and confirmatory laboratory investigations. Clinical criteria included fever (>38°C) or hypothermia (<36°C), tachycardia, dyspnea, assessment of general condition, and Apgar score. Laboratory investigations included blood culture, complete blood count, and C-reactive protein (CRP) measurement. Relevant maternal risk factors included discolored amniotic fluid, prolonged rupture of membranes, and vaginal infection during the third trimester. The exclusion criteria were as follows: Newborns whose parents or legal guardians did not provide informed consent; Newborns with congenital malformations; Newborns for whom no bacteriological samples were collected. Healthy newborns with negative discharge test results were included as controls.

**Table1:**
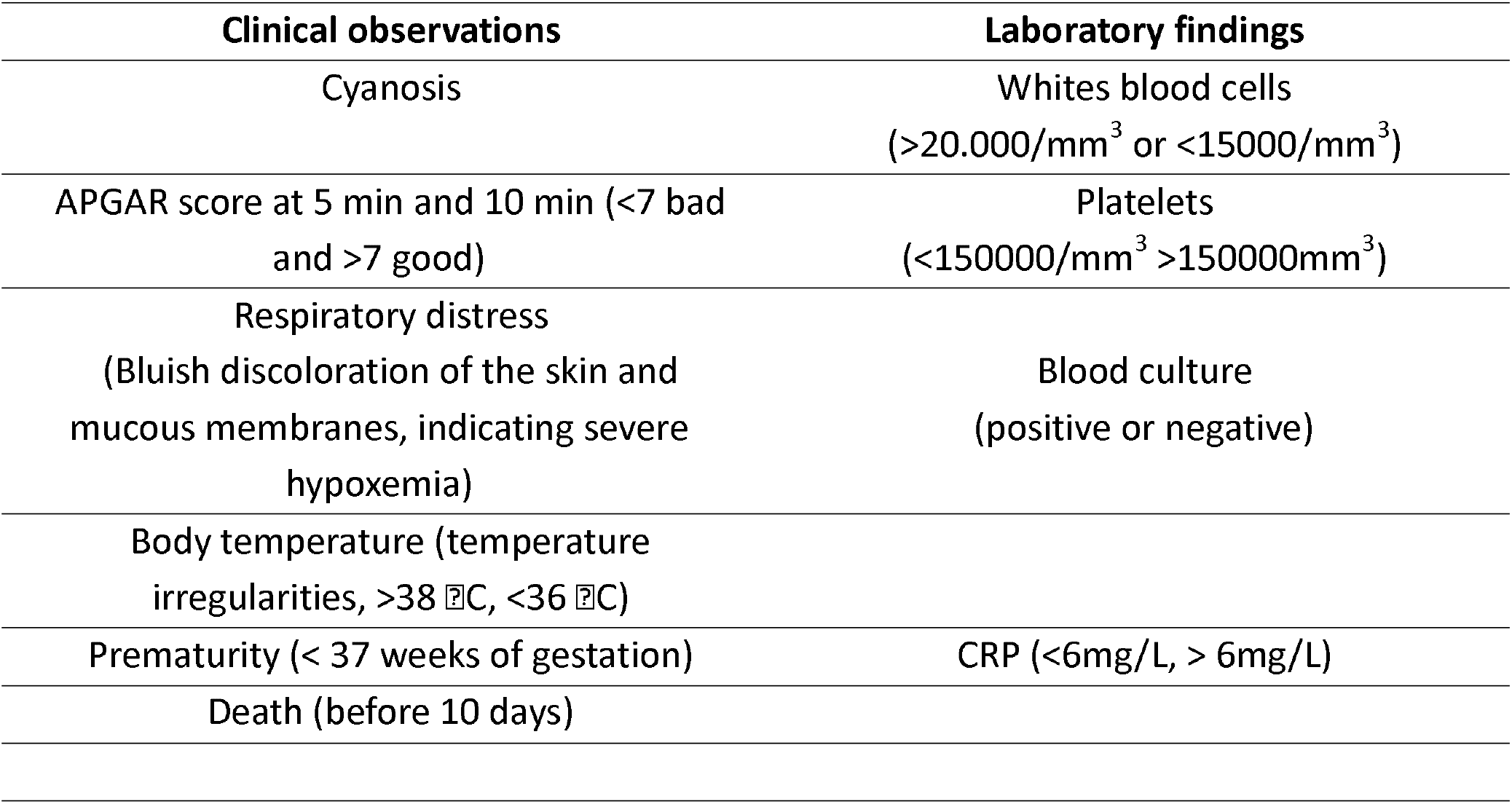
Clinical and biological parameters collected in the study. (APGAR stands for Appearance, Pulse, Grimace response, Activity, Respiration)

**Table2:**
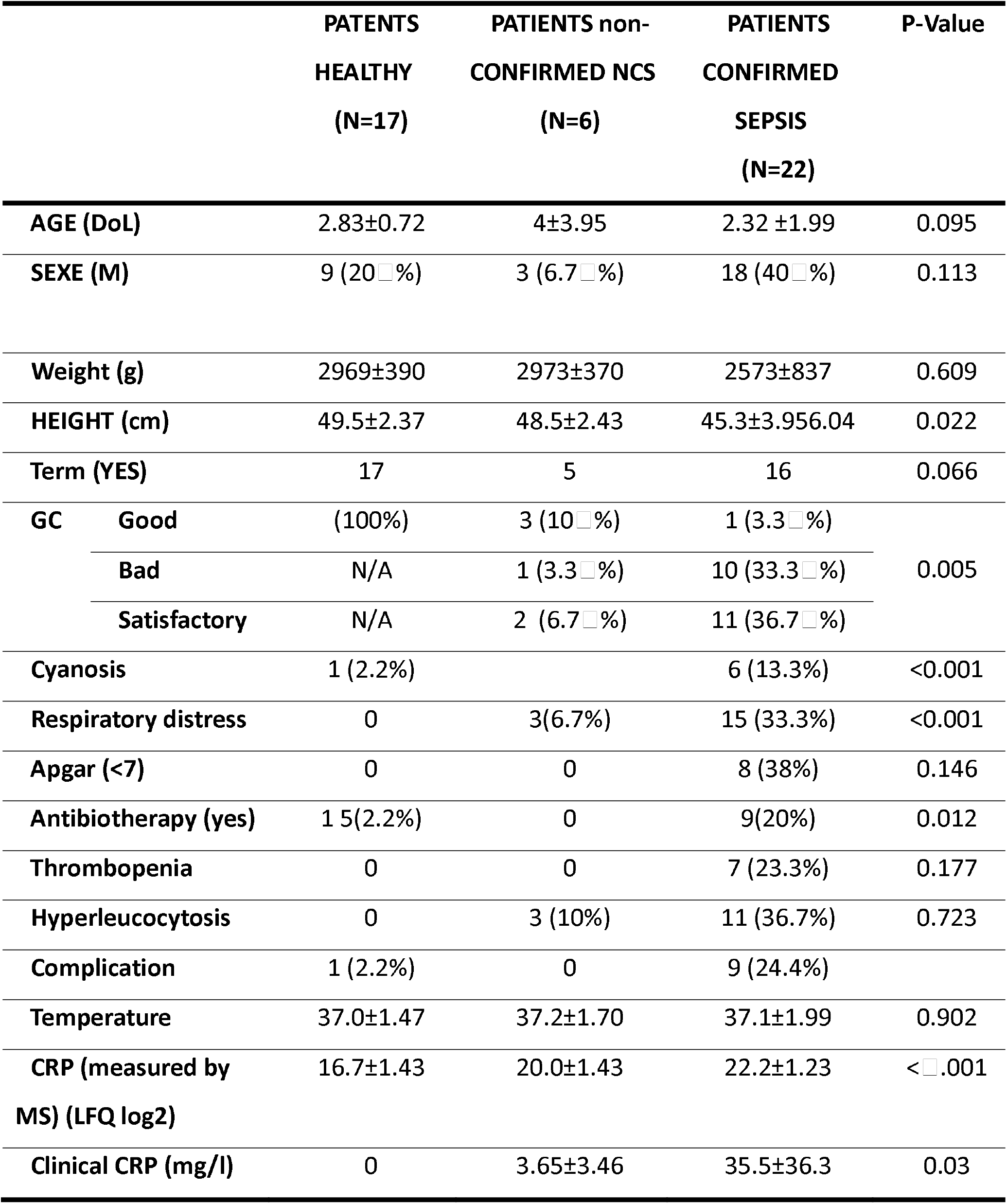
Descriptive statistics of patients clinical data collected in the study. (DoL = Day of Life ; M = Male ; GC = General Condition, CRP = C-Reactive Protein ; MS = Mass Spectrometry ; LFQ = Label Free Quantification)

Among the cohort of 60 newborns, four Healthy Control (HC) newborns were excluded because they had abnormal high CRP levels, with no further clinical information to evaluate their health state. Suspected but Non-Confirmed Sepsis (NCS) newborns were defined by the presence of at least two of the three following manifestations: hypothermia or hyperthermia; clinical signs of distress (respiratory distress, low APGAR score (Appearance, Pulse, Grimace response, Activity, Respiration); hematological abnormalities (including platelet and leukocyte counts outside the normal reference ranges). Confirmed sepsis (CS) cases were defined as patients with both positive blood culture and elevated C-reactive protein (CRP) intensity (as measured by mass spectrometry, exceeding 0 after z-scored of the log_2_ transformed LFQ intensity values). Additionally, CS had to present at least one of the three following criteria: thrombocytopenia; death before day 10 of life; prematurity. Six newborns presented contradictory profile and did not fall into these categories, were labelled as “Unclassified” and they were retained for unsupervised analysis (Supplementary Table 1) and 9 newborn were considered as Outlier based on clinical and/or analytical consideration.

Taking into consideration those criteria among the cohort of 60 newborns, 22 newborns were classified as CS, 6 newborns were suspected of infection and were therefore classified as NCS, 17 newborns were considered as HC, 6 could not be classified (Unclassified) finally 9 newborns were considered as Outliers.

The median DOL of the sampling was of 2.8 ± 0.72 for the heathy controls (HC), 4.0 ± 3.95 for the Non-Confirmed Sepsis (NCS) patients and 2.3 ± 1.99 for the Confirmed Sepsis patients (CS) (Table 2).

### Sample collection

Blood samples were collected in the neonatal unit of the pediatric department. Two samples were taken from the newborns, one on dry tube for serum and one with EDTA for plasma. Dry tube was immediately centrifuged at 3000 rpm for 5 minutes, the CRP was immediately measured on the serum with Architect I1000 automate by immunoturbidimetry method, while the rest of the serum samples were stored at -80°C and then transported in dry ice to the Proteomic Platform of Necker (Paris). EDTA tube was used for complete blood count.

### Proteomics sample preparation

S-Trap™ micro spin column (Protifi, Hutington, USA) digestion was performed on 150µg of neat serum according to manufacturer’s instructions. Briefly, samples were supplemented with 20% SDS to a final concentration of 5%, reduced with 20mM TCEP (Tris(2-carboxyethyl) phosphine hydrochloride) and alkylated with 50mM CAA (chloracetamide) for 5min at 95°C. Aqueous phosphoric acid was then added to a final concentration of 2.5% following by the addition of S-Trap binding buffer (90% aqueous methanol, 100mM TEAB, pH7.1). Mixtures were then loaded on S-Trap columns. Five washes were performed for thorough SDS elimination. Samples were digested with 2.5µg of trypsin (Promega) at 47°C for 1h. After elution, peptides were vacuum dried and resuspended in 2% ACN, 0.1% formic acid in HPLC-grade water prior to MS analysis.

### nanoLC-MS/MS analysis

Peptides were resuspended in 2% ACN, 0.1% formic acid in HPLC-grade water and 600ng were injected on an Evosep One system coupled to a timsTOF HT (Bruker Daltonics, Germany) mass spectrometer. The Evosep One system operated with the Whisper Zoom 40 Samples Per Day method using a 15 cm C18 Aurora Elite column (AUR3-15075C18-CSI, IonOpticks). The mobile phases comprised 0.1% FA as solution A and 0.1% FA/99.9% ACN as solution B. Mass-spectrometric data were acquired using the parallel accumulation serial fragmentation (PASEF) acquisition method in DIA (Data independent Analysis) mode with a 19 windows method using 33 Da windows covering the mobility ranges over a 400-1027 m/z range. The range of ion mobilities values from 0.67 to 1.29V s/cm2 (1/k0). The total cycle time was set to 1.06s.

### Data processing

Data analysis was performed using Spectronaut software (version 20.4). A search against the Homo sapiens (UniProtKB/Swiss-Prot) database (downloaded on 05 January 2026, 20420 entries) was performed using the directDIA+ (Deep) workflow with default settings.

Bioinformatic analysis was performed using R (v4.5.1) and Rstudio (RStudio 2025.09.1). All the preprocessing and statistical analysis was performed using proteoDA R package (v1.0.1) (Byrum, 2022/2026). For statistical comparison, raw Label Free Quantification (LFQ) intensities were log_2_-transformed, and we set five groups corresponding to the experimental conditions: 17 Healthy Control (HC), 6 Non-Confirmed Sepsis (NCS) 22 Confirmed Sepsis (CS), 6 unclassified and 9 outliers. The samples were labelled as outliers or unclassified according to inconsistency in clinical data and/or after exploratory data analysis of the experiment. The sample labelled as outliers were removed from the analysis whereas unclassified samples were kept in the analysis to investigate and compare their proteomic profile to the other samples. We then filtered the data to keep only proteins with valid values in at least 50% of the replicates in at least one experimental group. Next, missing values were imputed to fill missing data points by creating a Gaussian distribution of random numbers with a standard deviation of 33% relative to the standard deviation of the measured values and 1.8 standard deviation downshift of the mean to simulate the distribution of low signal values. Finally, limma-based statistics were performed between HC and NCS and CS groups with logFC threshold set a -0.5 and 0.5 and Benjamini-Hochberg multiple test correction was applied at 5% to control false discovery rate. Regarding the comparison between adult and newborn serum, all data analysis was performed using the same workflow and parameters except that no outliers were from the analysis and with logFC threshold set a -1 and 1.

Gene ontology over representation analysis (GO ORA) was performed on the significant dysregulated proteins using ClusterProfiler (version 4.18.4) and the ReactomePA (v1.54.0) R packages with org.Hs.eg.db R package (version 3.22.0) as a genome wide annotation for Homo sapiens with following arguments (pAdjustMethod = “BH”, pvalueCutoff = 0.05, qvalueCutoff = 0.05). Finally, enrichplot R package (version 1.30.4) was used for data visualization.

Plasma contaminant list was downloaded from the Quality Control of the Plasma Proteome Mannlabs github (https://github.com/MannLabs/Quality-Control-of-the-Plasma-Proteome/tree/master/data) and the specific contaminant markers from platelet were mapped in our adult/newborn dataset to calculate the platelet contaminant indexed for each sample by calculating the ratio of the LFQ intensities of the platelet markers against the sum of the other proteins quantified in our dataset.

Weighted gene co-expression network analysis (WGCNA) was performed on the normalized and imputed proteomics dataset using the R package WGCNA (v1.74). Protein abundance data were transposed such that samples were represented in rows and proteins in columns. Quality control was performed using the goodSamplesGenes function to identify potential outlier samples and proteins.

Then, a signed co-expression network was constructed and an optimal soft-thresholding power was determined using the pickSoftThreshold function (network Type set to « signed » and power ranging from 1 to 50) by assessing scale-free topology fit and mean connectivity across candidate powers. A soft-thresholding power of 10 was selected for network construction. Modules of co-expressed proteins were identified using the blockwise Modules function with a minimum module size of 10 proteins and module merging threshold of 0.25.

Module eigengenes were correlated with clinical traits, including hemoculture status, prematurity using Pearson correlation analysis and module–trait relationships were visualized using heatmaps with the ComplexHeatmap R package (v2.26.0).

## Results

### Neonatal Serum Proteome Characterization

Mass spectrometry-based proteomic analysis of neat serum revealed a significantly greater depth of coverage in 20 neonatal sera compared to 20 adult sera, with 979± 20 proteins identified on average in newborns versus 718± 40 in adults (p < 0.05). To investigate the basis for this enhanced detection, we examined the distribution of protein intensity through ranking plots comparing neonatal and adult healthy control samples (Figure 1A). Albumin concentration did not show any significative decrease in newborns, so it cannot account for the substantial increase in protein identifications (Figure 1B). In contrast, the top 50 most abundant proteins in adult serum, mostly immunoglobulins, are slightly lower in the newborn (Figure 1A, 1B). Platelet contamination, often correlated with higher protein identification^21^, was low and homogeneous among samples (Supplementary Figure 1A). The analysis revealed a slightly less compressed dynamic range in neonatal serum, with the curve displaced toward low abundance proteins. The expanded detection therefore likely reflects the developmental remodeling of immune system related protein expression patterns in early life. Hemoglobin isoforms exhibited expression profiles consistent with the expected perinatal transition from predominant fetal hemoglobin (HbF, α2γ2; encoded by HBA1/HBA2 and HBG1/HBG2) during the first 1–3 days of life to adult hemoglobin (HbA, α2β2; encoded by HBA1/HBA2 and HBB). Accordingly, fetal γ-globin chains (HBG1/HBG2) were more abundant and adult β-globin chains (HBB) less abundant in the first days of life, whereas HbA was predominant in adults, as previously described^22^ (Figure 1C). Very rare, low abundance fetal chains (HBZ, HBE1, HBM) were identified almost exclusively in some the newborns serum. This age-dependent hemoglobin switching served as an internal biological control, validating the capacity of our approach to capture known developmental transitions^8^.

**Figure 1:**
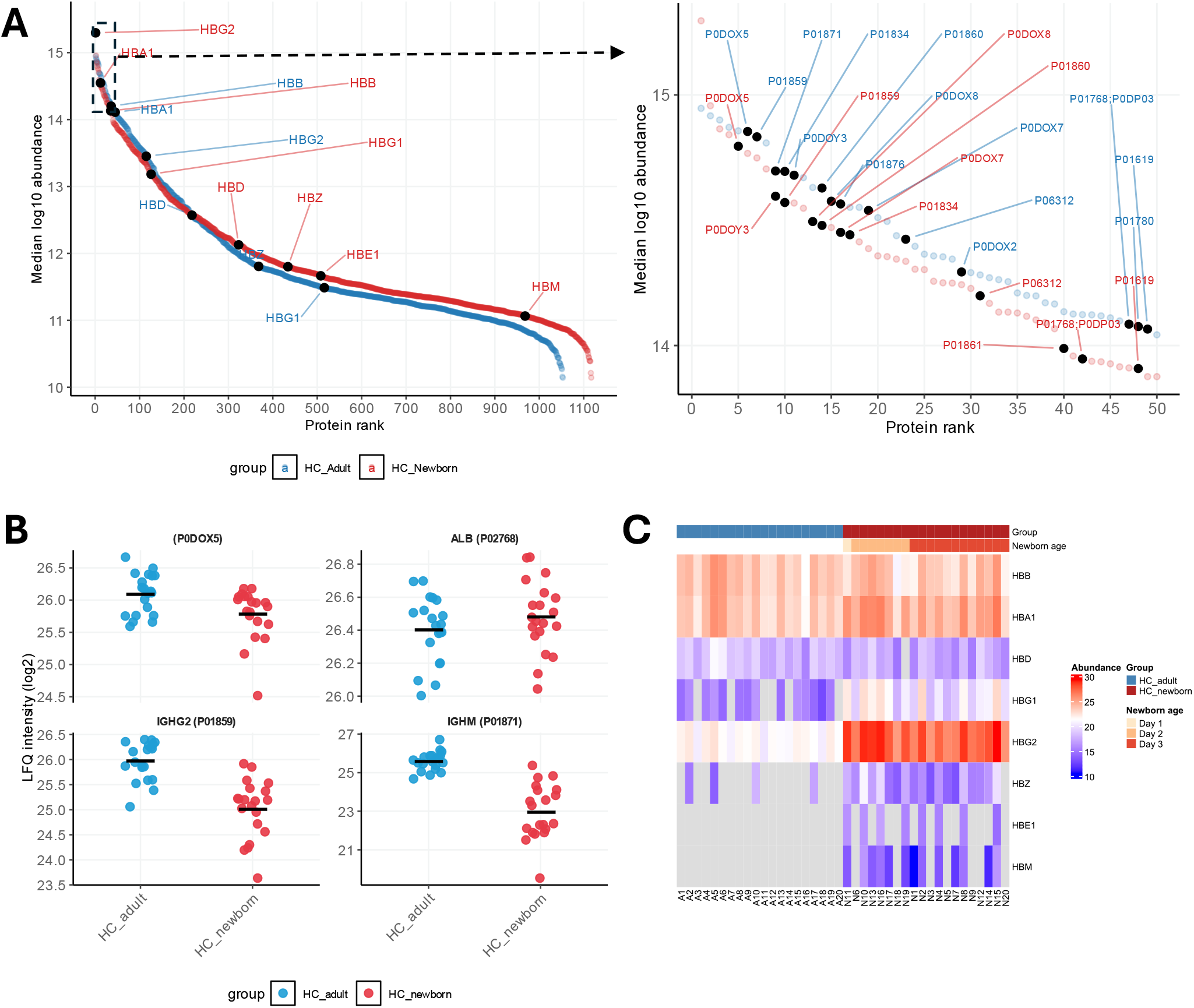
Comparison of proteome distribution and composition between healthy control adults and
newborns. (A) Ranking plots representing the comparison of protein distribution in Healthy Control (HC) adults vs HC newborns (left panel) along with a focus on the top50 proteins (right panel). (B) Profile plot of Albumin (ALB) and immunoglobulins (P0D0×5, IGHG2, IGHM. (C) Hemoglobin isoforms expression profiles in HC adults and HC newborns.

Comparison of neonatal and adult serum proteomes confirmed previously reported age-related differences and revealed novel protein classes enriched in the neonatal period. Principal component analysis (PCA) performed on 100% valid values separated neonatal from adult serum, indicating a distinct quantitative protein profile independently of data imputation (Figure 2A). Statistical comparisons identified 490 proteins significantly overrepresented in neonatal serum and 60 proteins overrepresented in adult serum (Figure 2B, Supplementary Table2). Additionally, 28 proteins were uniquely identified in neonatal samples, all of medium and low abundance as shown in the ranking plot (Supplementary Figure 1B, 1C).

**Figure 2:**
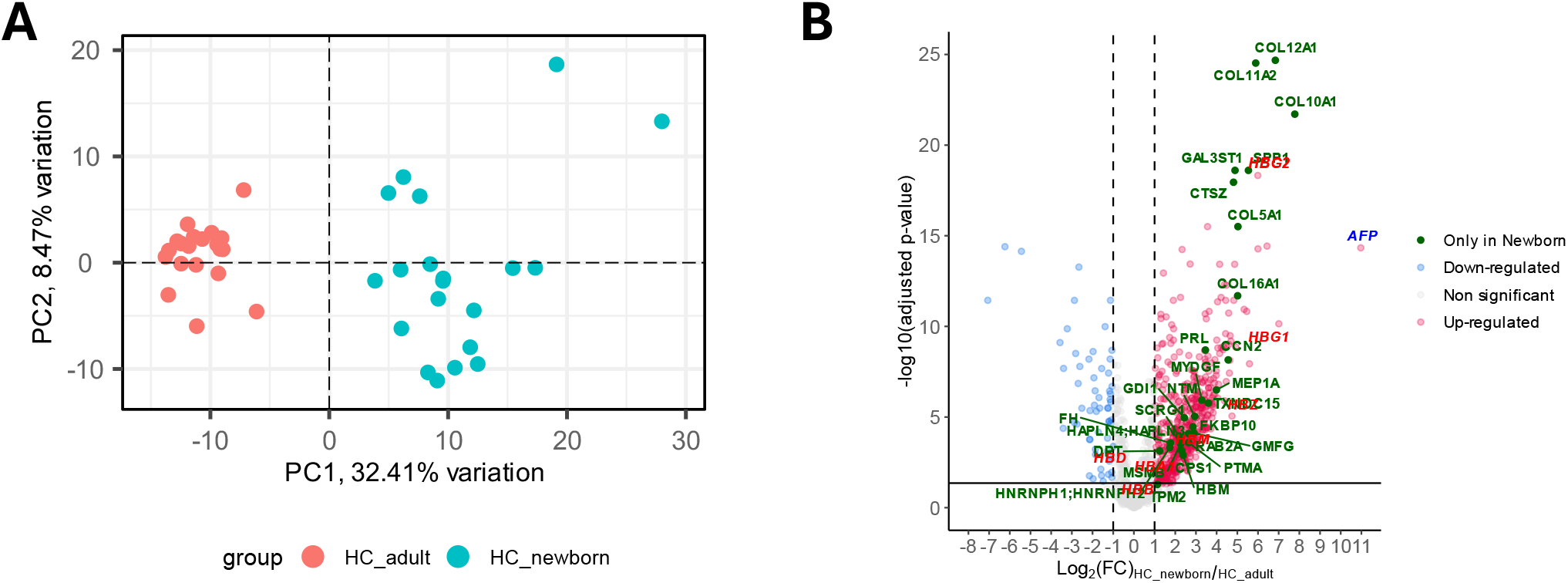
Clear separation of serum proteome from healthy control adults and newborns. (A) Principal Component Analysis performed with 100% valid values matrix (424 proteins over 1035 total proteins) on the 20 HC adults and 20 HC newborn. (B) Volcano plot of the statistical comparison HC newborn versus HC adult (490 proteins were found upregulated (red), 60 downregulated (light blue), 457 non significant (light grey), 28 found only in newborn (green) and none found only in adult.

In neonates compared to adults, consistent with prior studies, we observed elevated levels of proteins related to coagulation: α2-macroglobulin (P01023;A2M), periostin (Q15063;POSTN) and fibrinogen chains (P02671/P02675/P02679; FGA/FGB/FGG), while carboxypeptidase B2 (Q96IY4;CPB2) and heparin cofactor II (P05546; SERPIND1), were more abundant in adults^8,9^. We also observed a lower abundance in neonates of proteins related to hemoglobin regulation like haptoglobin (P00738;HP), haptoglobin-related protein (P00739;HPR), clusterin (P10909;CLU), and hemopexin (P02790;HPX), like in previous reports^8,9^ (Supplementary Figure 2). An extremely higher abundance of 2^10^ folds of alpha-fetoprotein was also confirmed in newborn serum^8^ (Figure 2B).

Functional enrichment analysis of proteins differentially abundant between neonates and adults revealed several overrepresented biological pathways. The most significantly enriched functional classes included immune response, in particular neutrophil degranulation, platelet activation and antigen processing for cross presentation. Reactome terms associated with metabolism of carbohydrates were also significantly overrepresented in the neonatal-enriched protein set for the first time (Figure 3A, 3B, Supplementary Table 3). Notably, we identified a new category of proteins significantly enriched in neonatal serum: collagen-related proteins and ECM, including 16 collagens, with the most abundant being COL16A1, COL12A1, COL10A1, COL11A2 (Figure 3A, Figure 2B).

**Figure 3:**
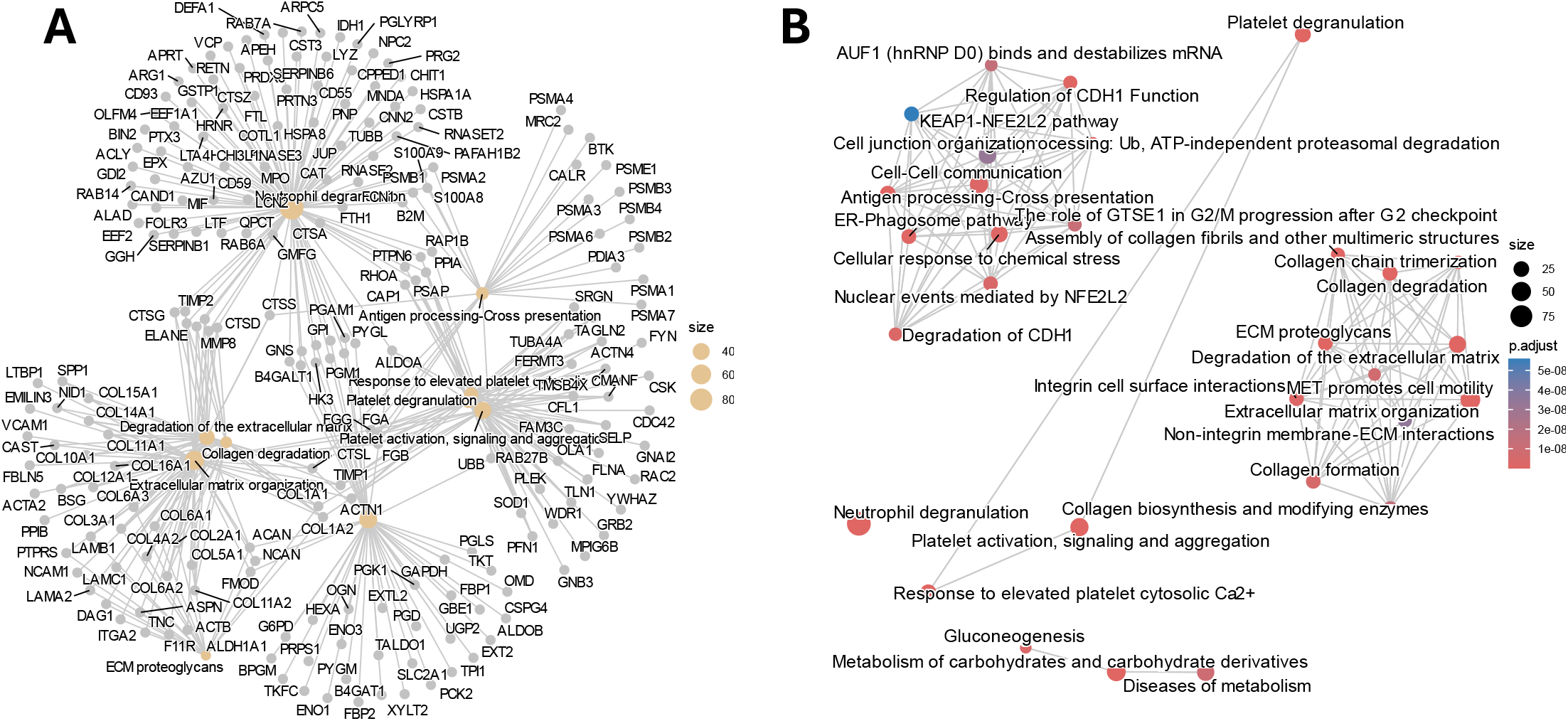
Over representation analysis from significantly upregulated proteins in HC newborns against HC adults. (A) Gene-concept network of the top 10 most significant Reactome pathways found after over representation analysis of the up-regulated proteins found in the HC newborn versus HC adult comparison (Figure 2B). (B) Enrichment map of the top 30 most significant Reactome pathways found after over representation analysis of the upregulated proteins found in the HC newborn versus HC adult comparison (Figure 2B).

Underrepresented classes of proteins in newborn include, coagulation factors including fibrinogen subunits and complement cascade; heme scavengers and oxygen transportation (Figure sup 3, Supplementary Table3), all in line with previous reports^8,9^.

### Sepsis Biomarker proteins

Statistical analysis between the three groups, defined at best of our knowledge as healthy control (HC), non-confirmed sepsis (NCS), confirmed sepsis (CS), provided a panel of 81 proteins significantly differential between CS and HC, with no significative protein between CS and NCS (Supplementary table 4). PCA analysis shows an influence of preterm/term status (PC1) and a separation on PC2 associated with positive hemoculture and confirmed sepsis (Supplementary Figure 4).

Given the heterogeneity of patient profiles and the inherent uncertainty in a posteriori diagnosis of confirmed sepsis, we performed WGCNA on the full proteomic dataset of 51 retained newborns in order to minimize potential bias due to misclassification. After hierarchical clustering of proteins based on co-expression pattern, 10 co-expression modules were identified through topological overlap-based dissimilarity after merging of modules highly correlated (Figure 4A, Supplementary Table 5). Then, correlation analysis between module eigengenes identified with WGCNA and clinical trait was performed in order to see positive or negative association. Among them, two clusters of strongly coregulated proteins were of particular interest: a cluster of 33 proteins (MEbrown in Figure 4B) that positively correlates with positive hemoculture, and a cluster of 20 proteins (MEred in Figure 4B) that negatively correlates with positive hemoculture. Moreover, the brown cluster showed a slight negative correlation with gestational term as well.

**Figure 4:**
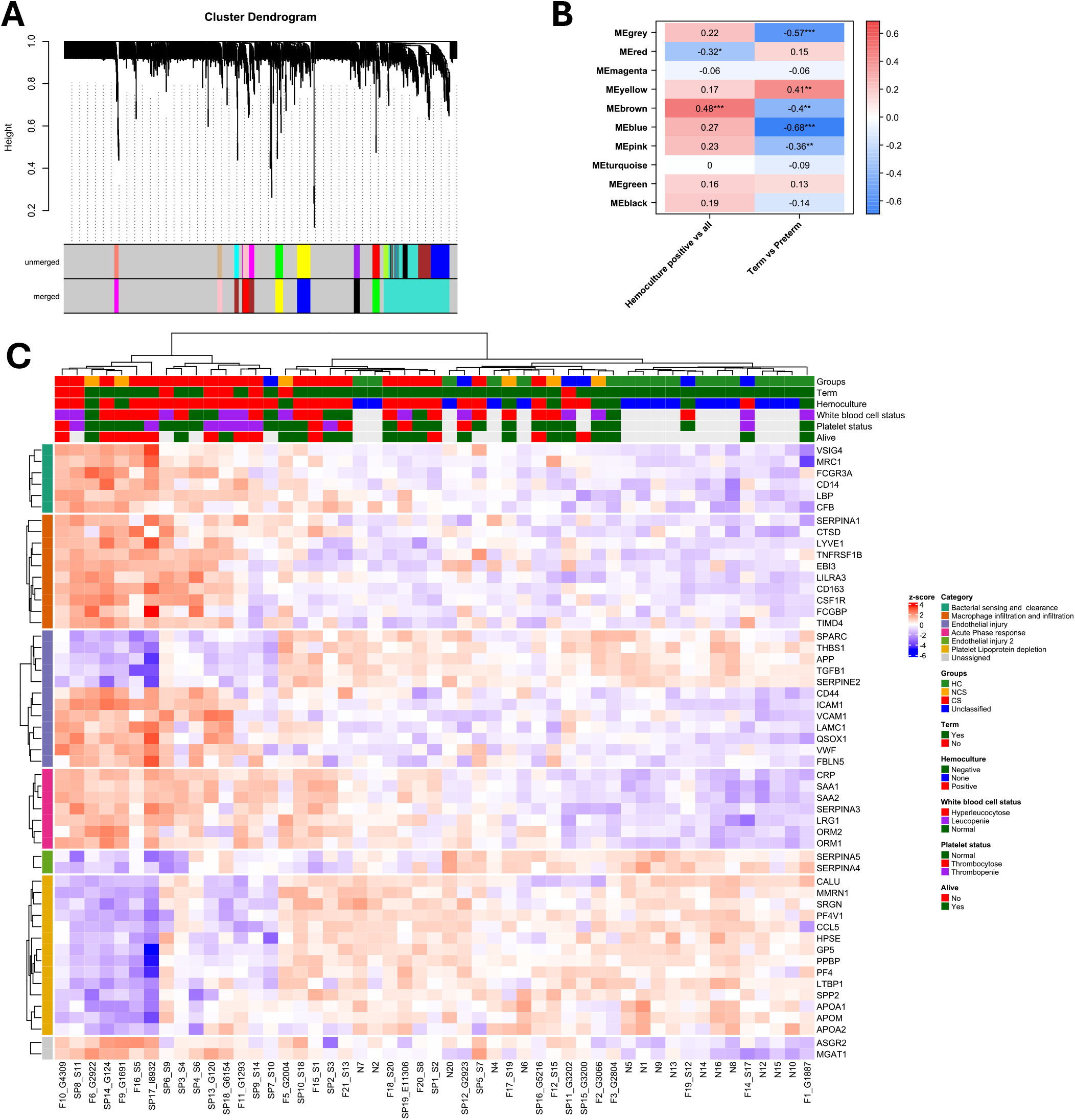
Weighted Gene Co-expression Network Analysis. (A) Hierarchical clustering dendrogram of proteins generated by weighted gene co-expression network analysis (WGCNA). Branches represent proteins clustered according to topological overlap–based dissimilarity. Module assignments identified by dynamic tree cutting before merging (“unmerged”) and after merging highly correlated modules (“merged”) are shown below the dendrogram as color bars. Each color corresponds to a distinct co-expression module. The merged module assignments were used for downstream analyses. (B) Correlation heatmap showing associations between module eigengenes identified by weighted gene co-expression network analysis (WGCNA) and clinical traits. Each row represents a co-expression module eigengene, and each column represents a clinical variable, including hemoculture positivity status and term versus preterm birth. Cell colors indicate the direction and strength of Pearson correlations, ranging from negative (blue) to positive (red), with white representing no correlation. Correlation coefficients and corresponding significance levels are displayed within each cell. (C) Heatmap representing the proteins belonging to the red and brown modules that significantly correlated positively (brown module) and negatively (red module) with hemoculture positivity status.

These two most relevant clusters were combined into a panel of 53 proteins and their expression profiles were plotted after z-score transformation (Figure 4C). This panel of 53 proteins successfully clustered patients in five sub-groups. Three sub-groups included 77% of hemoculture-positive cases and 91% of clinically classified CS cases. Amongst the proteins separating the patients, 29 were increased and 21 where underrepresented in these three subgroups. The fourth and fifth subgroups of patients were very similar in terms of protein intensity profile and regrouped 88% of the newborns classifies as HC.

This panel of 53 proteins contained proteins already reported in different studies as well as novel proteins highly correlated with them. LBP and CD14, both involved in endotoxin-recognition pathway, were increased. Furtermore, in our dataset the sequence coverage of CD14 is compatible with the detection of the secreted protein, also called presepsin (Supplementary Figure 5). We found increased CFB, VSIG4, FCGR3A, and MRC1 (CD206), consistent with opsonophagocytosis, macrophage-mediated bacterial clearance, and recognition of microbial glycans^23–25^. The strong elevation of CRP, SAA1, SAA2, ORM1, ORM2, SERPINA1, SERPINA3, and LRG1 indicates a robust hepatic acute-phase response, a known hallmark of severe infection and sepsis. Increased CD163, CSF1R, and TNFRSF1B further suggests monocyte/macrophage activation, and TNF-related inflammatory signaling compatible with sepsis-associated innate immune reprogramming^26^.

In parallel, we observe upregulation of ICAM1, VCAM1, vWF, CD44 all involved in endothelial dysfunction, and already reported as markers of severe sepsis in some studies (J. He et al., 2024). Conversely, deregulation and relative reduction of SERPINA4 and SERPINA5 was observed. This may reflect concomitant loss of vascular-protective, anti-inflammatory, and coagulation-regulatory mechanisms, consistent with sepsis-associated endothelial damage and coagulation imbalance^27^. Additional proteins including CD44, LYVE1, LAMC1, FBLN5, and QSOX1 suggested concomitant extracellular matrix remodeling and tissue repair processes, compatible with systemic inflammatory tissue injury occurring during severe sepsis^28,29^.

A striking feature of the downregulated proteomic profile was the coordinated reduction of multiple platelet alpha-granule including PF4, PPBP, CCL5, GP5, MMRN1, THBS1, SPARC, and SRGN, strongly suggesting platelet activation-associated granule depletion and consumptive coagulopathy. Simultaneous downregulation of APOA1, APOA2, and APOM further supports collapse of vascular-protective HDL-associated pathways, endothelial barrier stabilization, and anti-inflammatory lipid transport^30^.

Together, these findings support the detection of proteins specific to different processes involved in neonatal sepsis.

## Discussion

This work addresses the need of early and specific molecular protein markers to detect sepsis in newborns using unbiased, age specific, serum analysis by MS-based proteomics.

Relatively old proteomic studies have demonstrated that neonatal plasma has a distinct molecular composition compared to adults plasma^8,9^. To our knowledge, the dataset represented in this work is, the largest serum proteomic atlas currently available for healthy human newborns, with nearly 1,000 proteins consistently identified from neat serum. Our analysis confirmed several previously reported developmental differences between neonatal and adult circulation, including increased fetal hemoglobin chains, α2-macroglobulin, fibrinogen-associated proteins, and alpha-fetoprotein, together with lower abundance of heme-scavenging proteins such as haptoglobin and hemopexin^9,31,32^. Beyond validating known neonatal signatures, the depth of coverage enabled identification of additional biological programs that have not previously been characterized at the serum level in newborns. Most notably, we observed a striking enrichment of extracellular matrix and collagen-associated proteins, including multiple collagen isoforms and matrix remodeling proteins, suggesting that active tissue remodeling observed in skin may also be reflected in the neonatal circulating proteome^33,34^. Although COL1A1 has been reported before^8^, detection of a large family of ECM related in serum is unprecedented. In parallel, metabolic pathways linked to carbohydrate and glucose metabolism were significantly enriched, pointing to the profound metabolic adaptation during the immediate postnatal transition^35^. Together, these findings indicate that the neonatal serum proteome reflects not only immune immaturity and developmental hematopoiesis, but also intense systemic remodeling, growth, and metabolic reprogramming occurring during early extrauterine life.

With the same depth of analysis, we also investigated the serum proteome from neonates with suspected sepsis, with or without positive hemoculture, using an unsupervised, data-driven approach designed to capture latent biological pattern independently of imperfect clinical classifications and occasionally to record keeping. Rather than identifying a single static “sepsis signature”, the data suggests the existence of a dynamic continuum of partially overlapping host-response states in a subset of patients, which also correlated with positive hemoculture and clinical classification, although not entirely.

Despite the fact that many of the dysregulated proteins identified in our study have already been reported in independent studies using different approaches in neonatal and/or adult sepsis, the novelty of our work lies in the integration of these findings into a coherent framework linking the identified proteins to three major biological processes involved in sepsis: (i) pathogen sensing (LBP, CD14), and acute-phase inflammation (e.g. CRP, SAA1/2, ORM1/2); (ii) innate immune activation and leukocyte recruitment (e.g., FCGR3A, CSF1R, CD163, MRC1) and final platelet exhaustion and metabolic dysregulation, (e.g., PF4, PPBP, THBS1, GP5); (iii) endothelial injury and microvascular dysfunction with tissue remodeling (e.g., ICAM1, VCAM1, VWF, SPARC) accompanied by simultaneous loss of protective lipoproteins and serpins (e.g., APOA1, APOA2, APOM, SERPINA4, SERPINA5) (Figure 5).

**Figure 5:**
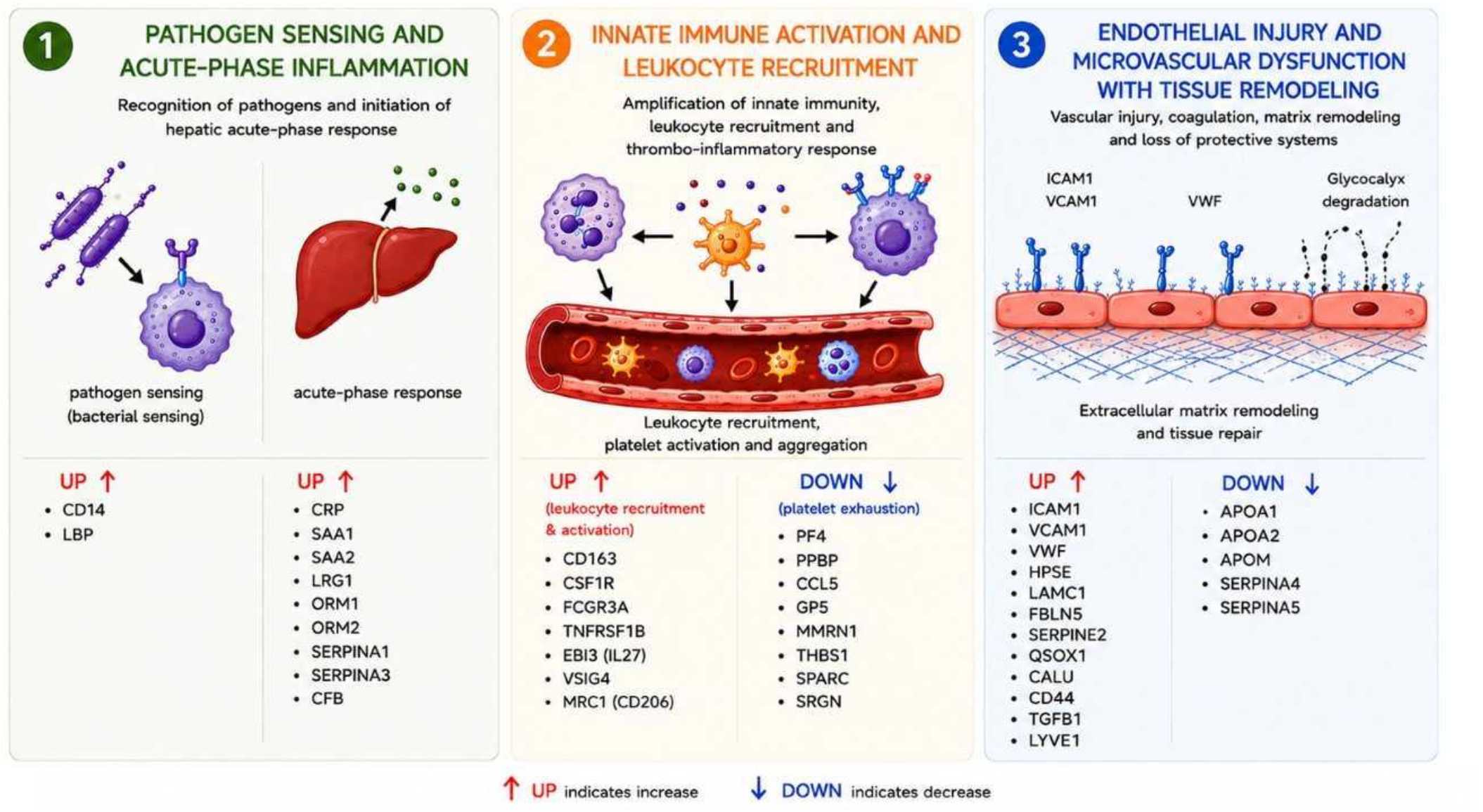
Summary of the three interconnected sepsis processes identified. Schema resuming the three interconnected sepsis processes identified. Proteins dysregulated in our study are associated to initial pathogen sensing and acute-phase response which triggers innate immune system activation and leukocyte recruitment, leading to platelet exhaustion and endothelial injury and repair attempts with loss of protective systems.

Pathogen-recognition molecules CD14 and LBP were increased, reflecting activation of the canonical endotoxin-recognition pathway, in which LBP facilitates transfer of bacterial lipopolysaccharide (LPS) to the CD14/TLR4 complex, a central mechanism in Gram-negative bacterial sepsis and systemic inflammatory activation^2,36,37^. Increased Presepsin, the soluble CD14-subtype, and is a well-documented putative biomarker of sepsis^38,39^, although it could also indicate chorioamnionitis^40^. The broadest detectable process was innate immune activation and hepatic acute-phase signaling. CRP, SAA1, SAA2, LRG1, ORM1, ORM2, SERPINA1, SERPINA3, and CFB were strongly elevated. However, these acute-phase reactant proteins — CRP, SAA1, SAA2, LRG1, ORM1, ORM2, SERPINA1, SERPINA3, and CFB — also can rise in several neonatal conditions beyond sepsis, including perinatal asphyxia^41^, meconium aspiration^42^, and even physiological stress of birth^43^. Although these proteins are not specific to sepsis, in some works a combination of them, such as CRP and SAA, have been found higher in sepsis than in non-infectious inflammation or controls in adults and neonates^44,45^. Of note, four of these proteins (LRG1, CRP, SAA1, SERPINA3) were also found and confirmed by immunodetection in a very recent study on umbilical cord of 16 in newborn with EOS^20^. Our larger pattern is compatible with rapid activation of IL-6–dependent acute-phase pathways following microbial exposure. Their concomitant detection with platelet, endothelial, and macrophage-remodeling pathways seen in this study appears more characteristic of culture-positive sepsis.

A distinct immunoregulatory program, from activation and recruitment of leukocytes to immune system shut down could also be observed. We observe increased TIMD4, LYVE1, CSF1R, CTSD, are all markers of resident/tissue macrophages^46^. Increase in proteins including CD163, MRC1 (CD206+), VSIG4, was consistent with activation of anti-inflammatory macrophage phenotypes associated with efferocytosis (CD163+, CD206+, VSIG4+) iron handling (CD163+, CD206+), and suppression of inflammatory injury and tissue-repair macrophages (CSF1R) often via IL-10 and TGF-β–dominated signaling^47^. Previous studies have shown that circulating sCD163 correlates with disease severity and organ failure scores and provides diagnostic and prognostic information beyond standard inflammatory markers such as CRP and procalcitonin^48,49^. Increased FCGR3A, TNFRSF1B and EBI3 (IL27) further suggested broad activation of innate myeloid programs. FCGR3A increase is in line with recent report of FCGR3A+ macrophage subset linked to prognosis in adult sepsis^50^.

Along with leukocyte recruitment, the proteomic signature indicates endothelial dysfunction and extracellular matrix remodeling. Elevated VWF, ICAM1, VCAM1, HPSE, LAMC1, SPARC, FBLN5, CD44 and FCGBP suggested activation of vascular inflammatory pathways associated with glycocalyx degradation, capillary leak, and microvascular injury^51^. ICAM1, VCAM1, VWF have already been reported as associated with negative outcome on adult sepsis patients^52^, although they can also be significantly elevated in patients following an acute ischemic stroke^53^. HPSE-mediated glycocalyx disruption has been implicated as a central driver of vascular permeability and organ dysfunction in sepsis, while increased VWF reflects endothelial activation and prothrombotic signaling^54^. These proteins were observed upregulated more strongly in one subgroup of patients, suggesting a later of more severe stage of the sepsis. Upregulation of QSOX1 and CALU may indicate activation of stress-response and extracellular matrix remodeling pathways accompanying vascular injury and tissue repair^55^.

The simultaneous reduction of platelet structural and granule-associated proteins, including PF4, PPBP, THBS1, GP5, MMRN1, and CCL5, further supports a model of consumptive platelet exhaustion accompanying endothelial injury and disseminated microvascular thrombosis^56^. In sepsis, sustained platelet activation promotes degranulation, immunothrombosis, and microvascular platelet consumption, ultimately leading to thrombocytopenia and exhaustion of circulating platelet granule components. In parallel, we observed depletion of circulating lipoproteins and vascular-protective mediators. APOA1, APOA2, APOM, SERPINA4, SERPINA5, were markedly reduced in the subclusters of patients associated with positive hemoculture. APOA1, APOA2, and APOM are key HDL-associated proteins involved in endotoxin buffering, endothelial stabilization, lipid transport, and anti-inflammatory signaling. In particular, ApoM-associated sphingosine-1-phosphate signaling is a critical regulator of endothelial integrity and vascular permeability during sepsis^57^. Their depletion has repeatedly been associated with severe sepsis and poor outcome^58^. Likewise, SERPINA4 and SERPINA5 regulate kallikrein, coagulation, and vascular permeability pathways, suggesting loss of endogenous vascular-protective buffering systems during septic progression. SERPINA4 has already been reported as a potential marker of sepsis, but not SERPINA5^59^. This coordinated collapse of lipoprotein-associated and anticoagulant protective programs may contribute directly to endothelial destabilization, uncontrolled thrombo-inflammation, and circulatory failure.

Importantly, these proteomic states did not segregate into rigid patient stages, but instead informed on a biological continuum spanning inflammatory activation, endothelial stress, thrombo-inflammatory amplification, and immunoregulatory vascular collapse. Three subclusters of patients showed deregulation of two or more of these biological processes (26 patients), including most — but not all — hemoculture-positive patients, whereas the remaining two subgroups (25 patients) included all healthy controls together with five previously unclassifiable patients. Amongst these five patients, SP11 and SP15 have been reported as hemoculture positive for S. Aureus, and with respiratory distress, but they presented low CRP, leukopenia and normal platelet count. According to their proteomic profile they cluster with non-confirmed sepsis. Clinical records for these patients are no longer available at the hospital, so further verification could not be conducted.

This multi-process marker panel may explain why isolated biomarkers such as CRP, single markers from other reprocesses (PPBP, SERPINA4, TIMD4, VCAM1, ICAM1) or platelet count alone often fail to robustly distinguish sterile inflammation from true sepsis in adults and neonates. The main limitation of this study is the relatively small number of patients included. However, these initial findings provide a valuable basis for replicating and validating the analysis in a larger, multicenter cohort. Such a study could facilitate the development of a comprehensive, condensed multi-process panel. Following feature reduction and evaluation of its robustness, it will open to the possibility of developing a multi-analyte immune test that could be used as a point-of-care in remote areas of Africa, where the access to hospital is limited. Indeed, quantification of the identified markers in leukocyte activation and platelet exhaustion could replace the platelet, leukocyte counting and CRP measurement when hospitals classic workups are not accessible. In addition, such panel will gain in specificity by including our proposed markers for vascular injury, leukocyte infiltration, pathogen sensing and clearing and multiple acute phase proteins.

Overall, our findings support the advantage to diagnose neonatal sepsis based on markers of multiple interconnected host-response processes. Larger multi-centric studies will be required to validate these proposed states and determine whether they can inform earlier diagnosis and therapeutic intervention.

## Acknowledgments

The authors are grateful to the parents of the patients who participated in this study. We thank all the members of the Proteomics Necker Platform (SFR Necker) for technical help. Artificial intelligence tools were used solely for language editing (spell checking and occasional rephrasing to improve English readability) and for improving the visual presentation of Figure 5. All outputs were critically reviewed and approved by the authors.

## Data Availability statement

The mass spectrometry proteomics data have been deposited to the ProteomeXchange Consortium via the PRIDE^60^ partner repository with the dataset identifier PXD079237. Reviewer access details: Log in to the PRIDE website using the following details: Project accession: PXD079237 Token: xxxxxxxxxxxx

## Ethics approval and consent to participate

The study was approved by the Research Ethics Committee of Cheikh Anta Diop University in Dakar. All procedures involving human participants were conducted in accordance with the ethical standards of the institutional research committee and the principles of the Declaration of Helsinki and its later amendments. Informed consent was obtained from all subjects and/or their legal guardian(s).

## Ethics declaration

The authors declare no competing interests.

Supplementary Figure 1: Platelet contamination profile and proteins exclusively found in healthy control newborns.

(a) Platelet contamination profile (top panel) and corresponding number of identified proteins (bottom panel) in each sample. Black line corresponds to the harmonic mean calculated from the platelet contaminant indexes of each sample and the red, orange and green dashed lines corresponding the 1, 2 or 3 standard deviation(s) from the harmonic mean, respectively.

(b) Ranking plot of the proteins only present in healthy control newborns (represented in black) after keeping proteins that have at least 50% of valid values in at least one of the experimental group.

(c) Heatmap representing the expression profile of the proteins only present in healthy control newborns after keeping proteins that have at least 50% of valid values in at least one of the experimental group.

Supplementary Figure 2: Profile plot representing the expression profile of the following proteins: A2M, AMBP, CLU, CPB2, FGA, FGB, FGG, HP, HPR, HPX, POSTN and SERPIND1 (gene name).

Supplementary Figure 3: Over representation analysis from significantly downregulated proteins in HC newborns against HC adults

(a) Gene-concept network of the top 10 most significant Reactome pathways found after over representation analysis of the downregulated proteins found in the HC newborn versus HC adult comparison (Figure 2B).

(b) Enrichment map of the top 30 most significant Reactome pathways found after over representation analysis of the downregulated proteins found in the HC newborn versus HC adult comparison (Figure 2B).

Supplementary Figure 4: Principal Component Analysis of the dataset from newborns suspected of sepsis.

(a) Principal Component Analysis performed with 100% valid values matrix (529 proteins over 1266 total proteins) on the 17 Healthy Control (HC), 6 Non Confirmed Sepsis (NCS) 22 Confirmed Sepsis (CS), 6 unclassified colored by experimental group.

(b) Principal Component Analysis performed with 100% valid values matrix (529 proteins over 1266 total proteins) on the 17 Healthy Control (HC), 6 Non Confirmed Sepsis (NCS) 22 Confirmed Sepsis (CS), 6 unclassified colored by term clinical trait.

Supplementary Figure 5: CD14 sequence coverage with peptides identified and quantified in proteomics

Supplementary Table1: Table of patient clinical data collected in the study

Supplementary Table2: Results of the HC adults against HC newborns statistical comparison

Supplementary Table 3: Results of the Over Representation Analysis performed on the significantly upregulated and downregulated protein of the HC adults against HC newborns statistical comparison

Supplementary Table 4: Results of the statistical analysis performed on the sepsis study

Supplementary Table 5: Results of Weighted Gene Co-expression Network Analysis performed on the sepsis dataset

