## Supplementary figures for "Serum proteomics in newborns reveals a multi-pathway molecular panel for neonatal sepsis, encompassing pathogen sensing, leukocyte recruitment, and endothelial injury"

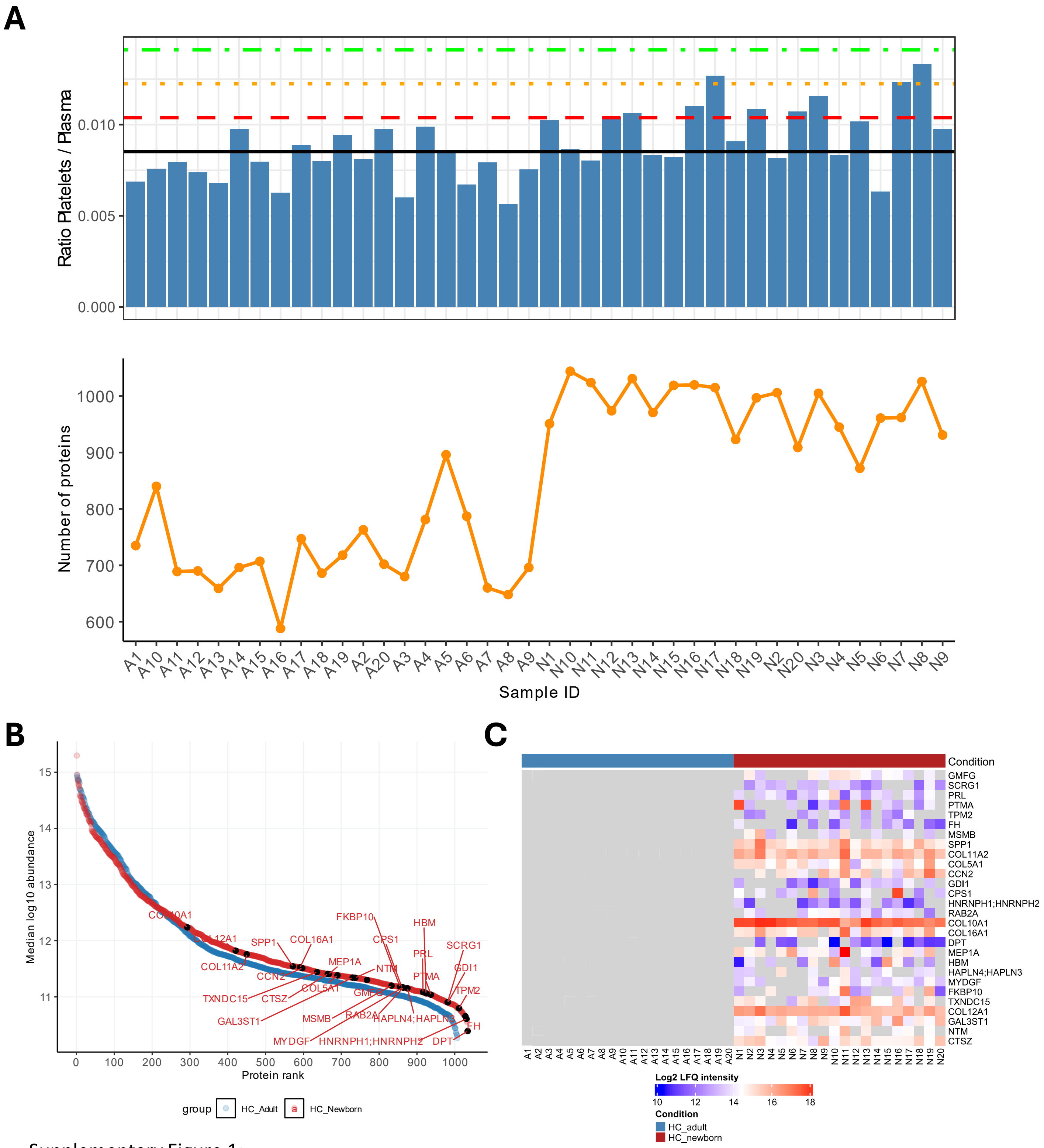


Supplementary Figure 1:

(A) Platelet contamination profile (top panel) and corresponding number of identified proteins (bottom panel) in each sample. Black line corresponds to the harmonic mean calculated from the platelet contaminant indexes of each sample and the red, orange and green dashed lines corresponding the 1, 2 or 3 standard deviation(s) from the harmonic mean, respectively.

(B) Ranking plot of the proteins only present in healthy control newborns (represented in black) after keeping proteins that have at least 50% of valid values in at least one of the experimental group

(C) Heatmap representing the expression profile of the proteins only present in healthy control newborns after keeping proteins that have at least 50% of valid values in at least one of the experimental group

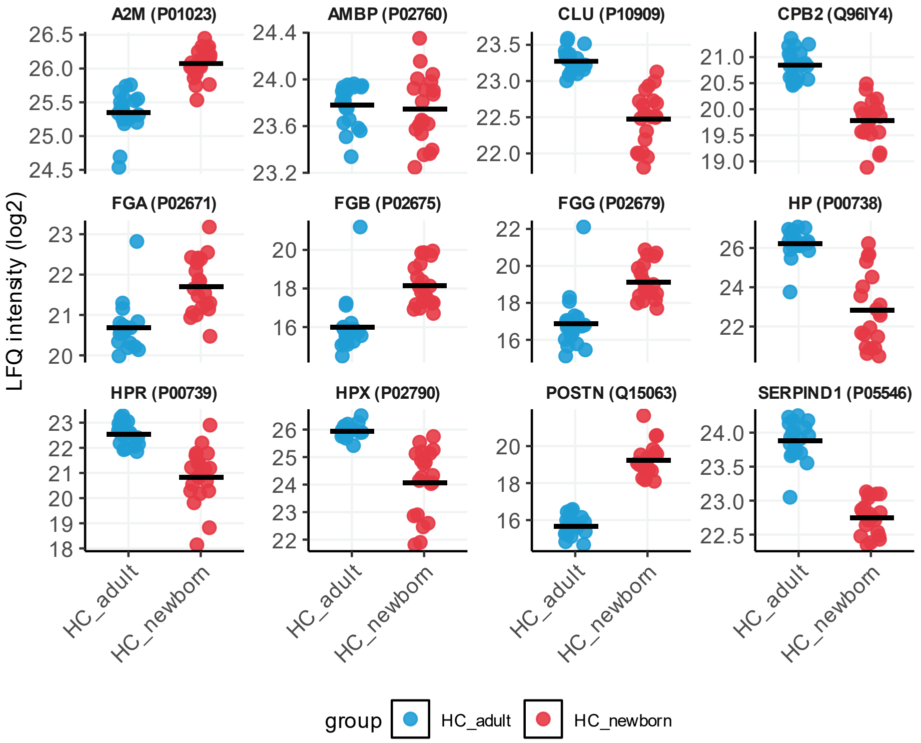


Supplementary Figure 2:
Profile plot representing the expression profile of the following proteins: A2M, AMBP, CLU, CPB2, FGA, FGB, FGG, HP, HPR, HPX, POSTN and SERPIND1 (gene name)

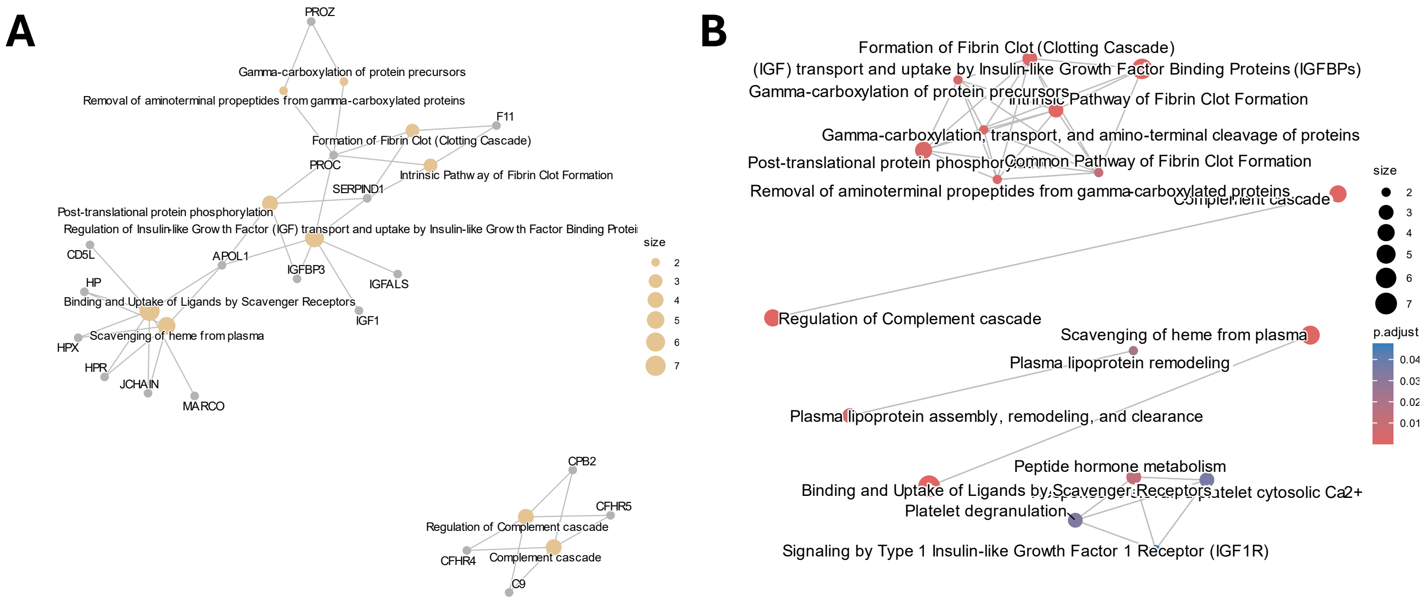


Supplementary Figure 3:

(A) Gene-concept network of the top 10 most significant Reactome pathways found after over representation analysis of the downregulated proteins found in the HC newborn versus HC adult comparison (Figure 2B)

(B) Enrichment map of the top 30 most significant Reactome pathways found after over representation analysis of the downregulated proteins found in the HC newborn versus HC adult comparison (Figure 2B)

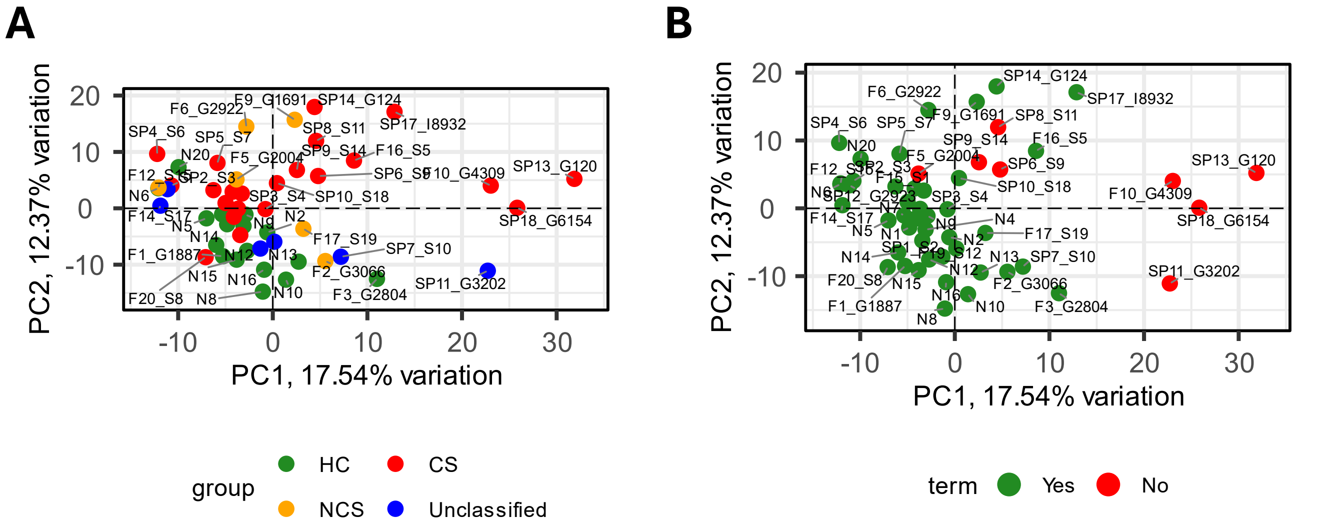


Supplementary Figure 4:
Principal Component Analysis performed with 100% valid values matrix (529 proteins over 1266 total proteins) on the 17 Healthy Control (HC), 6 Non Confirmed Sepsis (NCS) 22 Confirmed Sepsis (CS), 6 unclassified colored with experimental group (A) or term status (B)

>sp|P08571|CD14_HUMAN Monocyte differentiation antigen CD14 OS=Homo sapiens OX=9606 GN=CD14 PE=1 SV=2

MERASCLLLLLLPLVHVSATTPEPCELDDEDFRCVCNFSEPQPDWSEAFQCVSAVEVEIH
AGGLNLEPFLKRVDADADPRQYADTVKALRVRRLTVGAAQVPAQLLVGALRVLAYSRLKE
LTLEDLKITGTMPPLPLEATGLALSSLRLRNVSWATGRSWLAELQQWLKPGLKVLSIAQA
HSPAFSCEQVRAFPALTSLDLSDNPGLGERGLMAALCPHKFPAIQNLALRNTGMETPTGV
CAALAAAGVQPHSLDLSHNSLRATVNPSAPRCMWSSALNSLNLSFAGLEQVPKGLPAKLR
VLDLSCNRLNRAPQPDELPEVDNLTLDGNPFLVPGTALPHEGSMNSGVVPACARSTLSVG
VSGTLVLLQGARGFA

Supplementary Figure 5:
CD14 sequence coverage with peptides identified and quantified in proteomics
